# The D1 receptor agonist SKF38393 improves waiting impulsivity in a baseline dependent manner

**DOI:** 10.1101/2023.07.15.549161

**Authors:** Sara Abdulkader, John Gigg

## Abstract

**Rationale:** Stimulants are the first-line treatment for attention-deficit/hyperactivity disorder (ADHD). However, the ensuing risk of abuse with stimulants means there is an urgent need for new, low-risk therapeutic agents. D1 receptors play an important role in the cognitive enhancing effects of stimulants and thus may provide a therapeutic target. Previous pre-clinical studies have shown that selective activation of D1 receptors improves sustained attention in the 5C-CPT without improving waiting impulsivity (premature response).

**Objective:** The aim of the present experiment was to compare the effects of the selective D1 receptor agonist SKF 38393 to a standard ADHD treatment (amphetamine) on waiting impulsivity in the 5C-CPT under extended inter-trial intervals. Oldham’s method was used to determine the presence of a rate-dependent effect.

**Methods:** Adult female Lister hooded rats were trained to criterion in the 5C-CPT (>70% accuracy, < 30% omission and < 40% false alarms). Effects of the selective D1 receptor agonist SKF 38393 (2-6 mg/kg) or amphetamine (0.1-0.4 mg/kg) were investigated under behavioural manipulations to challenge inhibitory response control.

**Results:** The highest dose of SKF 38393 and the two highest doses of amphetamine improved waiting impulsivity in a baseline dependent manner. The clockwise movement of the regression line indicates that, as the dose increases, the magnitude of improvement increases in a manner consistent with baseline performance.

**Conclusions:** These findings support further clinical investigation of D1 receptor modulators to facilitate the discovery of improved medications for impulsive behaviour related disorders such as ADHD. The concept of rate dependency applies to effects of SKF 38393 or amphetamine on waiting impulsivity. Oldham’s correlation method may present an opportunity to enhance the translational value of research in the preclinical laboratory to the clinic.

## Introduction

Attention deficit hyperactivity disorder (ADHD) is a psychiatric condition characterized by three main behavioural deficits: impulsivity, hyperactivity and lack of attention (Mueller et al., 2017; Storebø et al., 2023). These are observed during early childhood and often persist into adult life. Indeed, children diagnosed with ADHD have a higher risk of developing social and cognitive problems in later life (Lara et al., 2009; Retz et al., 2021). The prevalence of ADHD is between 2-7%: 5% of children have difficulties with inattention and hyperactivity and 2-4% of adults are diagnosed with ADHD (Sayal et al., 2018). With increasing age, inattention becomes more prominent while impulsivity diminishes (Faraone et al., 2015; Telzer et al., 2016). Depending on the main symptoms, ADHD can be divided into three categories: lack of attention (most prevalent among girls), hyperactivity (more prevalent among boys) and a third subtype that is a combination of the first two (Rasmussen and Gillberg, 2000; Wu et al., 2022). ADHD is more common in boys than in girls (5.2% vs. 2.7%) (Sayal et al., 2018; Young et al., 2020; Mohammadi et al., 2021). ADHD is unrecognised, underdiagnosed and under-treated in the United Kingdom (Young et al., 2021). In the UK, less than 50% of children with inattention and hyperactivity have been diagnosed with ADHD (Faraone et al., 2015; Young et al., 2021). Girls may go underdiagnosed because they tend to have inattentive traits (less noticeable symptoms) rather than impulsive traits (Faraone et al., 2015; Mohammadi et al., 2021). In addition, co-existing anxiety and depression in girls with ADHD may lead to missed diagnoses (Quinn & Madhoo, 2014).

Stimulants such as methylphenidate (Ritalin) and amphetamine (Amfetamine) are the most common pharmacological treatments for ADHD (Mueller et al., 2017; Storebø et al., 2023). However, only 40% of patients respond to both or only one of these drugs (Wolraich et al., 2019). Short-term treatment with stimulants has been linked with headache, abdominal pain, reduced appetite (e.g., anorexia) and sleep disturbances. On the other hand, long-term treatment with stimulants have been linked with slower growth rate in children, for example, methylphenidate causes growth retardation of 1.38 cm/year (Pliszka et al., 2006: Hodgkins et al., 2012; Carucci et al., 2021). It has also been shown that stimulant medications increase the risk of cardiovascular diseases (Hennissen et al., 2017; Farrell et al., 2019). Importantly, stimulants have a high potential for abuse and dependence (Farrell et al., 2019; Shellenberg et al., 2020; Hersey et al., 2021). Amphetamine (AMPH) leads to dependence in 11% of those who use it (Farrell et al., 2019). Prescription stimulant medication misuse is a serious problem (Weyandt et al., 2016; Sharif et al., 2021). For example, stimulants (‘smart pills’) are commonly used by students to enhance academic performance (Sharma et al., 2014; De Bruyn et al., 2019). Misuse of prescription stimulants has dangerous side effects such as euphoria, hallucinations and aggression (Shellenberg et al., 2020; Hersey et al., 2021). Food and drug administration encourages the development of abuse-deterrent formulations for psychostimulants (Shellenberg et al., 2020). Non-stimulants (such as atomoxetine and guanfacine) are the second-line treatment for ADHD; however, these are less effective than stimulants and have a delayed onset of action. It has also been shown that atomoxetine and guanfacine negatively influence mesenchymal stem cell migration during bone formation (Wagener et al., 2022). Therefore, there is a pressing need for new therapeutic agents that act without these side effects (e.g., reduced physical growth) and dependence potential (Sharma et al., 2014).

One of the characteristic features of ADHD is impulsivity (behavioural disinhibition), which is important to minimise in order to suppress inappropriate action (Arnsten, 2009). Behavioural disinhibition may lead to actions being carried out rapidly, without thinking about their consequences, or it may lead to compulsivity, which refers to repetitive and persistent performance of a certain action even though it does not lead to a reward (Dalley et al., 2011; Faraone et al., 2015). Deficits in four subtypes of impulsivity have been reported in ADHD: (a) response disinhibition (impulsive action), which involves difficulty in inhibiting a pre-potent response; (b) poor waiting impulsivity, which is seen as difficulty in waiting for stimulus presentation before responding; (c) temporal discounting (decisional impulsivity or impulsive choice), which involves preference for immediate over delayed reinforcement; and (d) poor stopping impulsivity (action cancellation), which manifests as a difficulty in stopping an ongoing response (Dalley et al., 2011). Waiting impulsivity, typically measured in rodents using 5-choice serial reaction time task (5-CSRTT), is modulated by the mesolimbic dopaminergic system (reward pathway), which projects from the VTA to the limbic system (Dalley et al., 2011; Dalley and Robbins, 2017). The ventral striatum (particularly the nucleus accumbens (NAc)) has an important role in regulating waiting impulsivity (Dalley and Robbins, 2017), for example, poor performance of high-impulsive rats on the 5-CSRTT has been linked with a deficit in dopaminergic function within the NAc (Robinson et al., 2009). The 5-CSRTT was later modified to the 5C-CPT (Young et al., 2009; Barnes et al., 2012), which incorporates non-target trials where the rat must withhold responding at all five lit positions (Young et al., 2009; Tomlinson et al., 2014; Hayward et al., 2016). This incorporation of non-target trials helps to dissociate response inhibition from waiting impulsivity; premature responses reflect a deficit in waiting impulsivity, while the failure to withhold responding (false alarms) during non-target trials represents response disinhibition (Young et al., 2011; Tomlinson et al., 2014). This task also allows the assessment of vigilance in rodents in a manner consistent with human CPT using signal detection theory (Tomlinson et al., 2014). In addition, a variable ITI is incorporated in 5C-CPT to prevent rats from simply timing their response (Van Enkhuizen et al., 2014). D1 receptors are mainly expressed in the PFC and NAc, critical regions in the circuit mediating waiting impulsivity (Weiner et al., 1991; Vincent et al., 1993; Pezze et al., 2007; Sasamori et al., 2019). D1 receptors have long been implicated in attentional processes and inhibitory response control as measured by the 5-CSRTT and 5C-CPT (Barnes et al., 2012; Zhu et al., 2017).

Evidence collected by our group suggests that activation of specific dopamine receptor subtypes (particularly D1 and D4) can alter behavioural measures associated with 5C-CPT performance. For example, administration of a D1 receptor agonist (SKF-38393) enhanced vigilance by increasing hit rate without affecting response inhibition (the probability of false alarm) under challenging conditions (extended ITI-define) in rats (Barnes et al., 2012), suggesting that selective activation of D1 receptors represents a promising approach to the treatment of ADHD. However, this drug impaired performance under standard task conditions, suggesting that the effectiveness of D1 receptor agonists in enhancing attention depends on basal dopamine levels (Barnes et al., 2012). Thus, under high task demands, activation of D1 receptors improves certain domains of cognitive function (such as attention) without affecting other domains such as waiting impulsivity and response inhibition (Barnes et al., 2012). Despite these findings, the role of the D1 receptor on waiting impulsivity under an extended ITI is yet to be determined. It has been reported that activation of the dopaminergic system improves performance of low-performing animals but impairs performance of high-performing animals, suggesting that the relationship between attentional performance or inhibitory response control and dopamine level follows an inverted-U shaped curve (Tomlinson et al., 2015; Hayward et al., 2016; Bickle et al., 2016; Caballero-Puntiverio et al., 2020). These studies indicate that increasing dopamine neurotransmission under high task demands by activation of specific receptor subtypes normalises the dopaminergic deficit in low performing rats; however, further activation of a normal dopaminergic system in high performing rats will impair performance (Tomlinson et al., 2014; Hayward et al., 2016). Although there is evidence to support the baseline-dependent effect of stimulants on 5-CSRTT or 5C-CPT performance, few studies have used correlational analysis to explicitly test this.

Indirect dopaminergic agonists such as AMPH, methylphenidate and modafinil are widely used in the clinic to treat cognitive deficits (Mueller et al., 2017; Storebø et al., 2023). These drugs are also commonly used in preclinical studies to test the predictive validity of behavioural tasks for assessing pharmacologically induced enhancement in cognitive functions, particularly attention and impulsivity (Tomlinson et al., 2014; MacQueen et al., 2018). Recently, it has been reported that touchscreen 5C-CPT can reliably reveal a cognitive enhancing effect of AMPH in mice and human, as systemic administration of AMPH (a potent dopamine and noradrenaline reuptake inhibitor) under baseline (standard task demand) conditions improved vigilance by increasing hit rate without affecting response inhibition (probability of false alarm) and waiting impulsivity (premature response) in both species (MacQueen et al., 2018). In addition, clinical trials report that low-moderate doses of AMPH improves inhibitory response control in individuals with high baseline impulsivity (Arkell et al., 2022). Collectively, these findings support the role of the dopaminergic system in cognitive function, however, the exact mechanisms by which drugs that increase activity of the dopaminergic system improve cognitive function in individuals with ADHD is not fully understood (Arnsten, 2009; Mueller et al., 2017; Faraone et al., 2015). Understanding the mechanism of action of these drugs will facilitate drug discovery efforts for ADHD. Correlational analysis may present an opportunity to enhance the translational value of research in the preclinical laboratory to the clinic (Abdulkader et al., 2023). This method can provide significant insight into the neural mechanism underlying therapeutic effects of stimulants. Previous studies suggest that D1 receptor activation contributes to the therapeutic effects of psychostimulants (methylphenidate and modafinil) and atomoxetine (Arnsten and Dudley, 2005; Gamo et al., 2010; Yin et al., 2022). The aim of the present study was to examine the role of AMPH and a D1 receptor agonist on inhibitory response control as assessed by 5C-CPT performance. AMPH was used to test the predictive validity of 5C-CPT for assessing pharmacologically induced enhancement in waiting impulsivity. Previous studies have reported no change in inhibitory response control following administration of SKF-38393 under task conditions that target attention (Barnes et al., 2012). Therefore, we challenged animals with an extended ITI and session length to determine whether SKF 38393 can alter waiting impulsivity under such challenging task conditions that target inhibitory response control.

## Materials and Methods

### Animals

Adult female Lister hooded rats (n=15; Charles River, UK; Weighing 240 ± 10g at the start of the experiment) were housed in groups of 5 in a temperature (21 ± 2 °C) and humidity (55±5%) controlled environment (University of Manchester BSF facility). They were allowed free access to food (Special Diet Services, UK) and water for one week prior to the beginning of training. Home cages were individually ventilated cages with two levels (GR1800 Double-Decker Cage, Techniplast, UK) and testing was completed under a standard 12-hour light: dark cycle (lights on at 07:00 am). Two days before training commenced, food restriction was initiated to encourage their task engagement. This restriction continued throughout training where rats were maintained at approximately 90% of their free-feeding body weight (fed 10 g rat chow/rat/day) with free access to water. All procedures were conducted in accordance with the UK animals (Scientific Procedures) 1986 Act and local University ethical guidelines.

### Rat 5C-CPT Apparatus

Training took place in 8 operant chambers (Campden instruments Ltd, UK), each placed inside a ventilated and sound-attenuating box. The chambers were of aluminium construction 25cm × 25 cm. Each chamber had a curved wall containing nine apertures. For 5C-CPT training, four of these apertures were blocked by metal caps while the other five were left open (numbers 1, 3, 5, 7 and 9 were open). Each aperture had a light and an infrared beam crossing its entrance to record beam breaks due to nose poke. A reward dispenser was located outside each chamber that automatically dispensed reinforcers (45 mg sucrose food pellets; Rodent Pellet, Sandown Scientific, UK) into a food tray at the front of the chamber. Entrance to each food tray was covered by a hinged panel so that tray entries were recorded when the rat’s snout opened the panel. House lights were placed in the side wall of the chambers and performance was monitored by a camera attached to the chamber ceiling.

### 5C-CPT behavioural training

5C-CPT training was carried out 5 days per week. A session consisted of 120 trials or was 30 min long, whichever came first. The training consisted of 3 phases: (i) 5-CSRTT training, (ii) 5C-CPT training with fixed inter-trial interval (ITI) and (iii) 5C-CPT training with variable ITI (Tomlinson et al., 2014; 2015). Task parameters including stimulus duration and limited hold were gradually reduced for rats based on their individual performance.

### Signal detection theory (SDT)

Two primary parameters are calculated from 5C-CPT trial-outcome measures. The first is hit rate, which is the proportion of ‘hits’ to ‘misses’ and the second is false alarm rate, which is the proportion of false alarms to correct rejections (Young et al., 2011; Table 2). From these two parameters, sensitivity (vigilance) and bias can be calculated using SDT (Young et al., 2011). Sensitivity reflects the ability to respond when there is only one illuminated hole and inhibit that prepotent response when all five holes are illuminated. Sensitivity can be estimated by calculating the discriminability index (*d*’), which is the difference between z score of hit rate and false alarm rate (Young et al., 2009; Anderson, 2015). 5C-CPT performance can also be affected by bias (tendency to respond to stimuli), which can be measured non-parametrically by calculating the responsivity index (RI). A positive RI value reflects bias towards FA (liberal response strategy), whilst a negative RI reflects bias towards omission (conservative response strategy). When RI is zero there is no bias (Anderson, 2015; Young et al., 2013).

## Experimental Design

### Drug preparation

d-Amphetamine sulphate (Sigma, UK) was dissolved in 0.9 % saline. SKF 38393 a selective D1 receptor agonist (Dubois et al., 1986; Barnes et al., 2012), (Sigma, UK) was dissolved in distilled H_2_O. Both drugs were injected via the intraperitoneal (i.p.) route at a volume of 1 ml/kg. All drug doses were calculated as base equivalent weight. The doses used in this study were chosen based on previous studies performed in the same strain and sex of rats (Barnes et al., 2012). AMPH doses were selected to be lower than those that increased locomotor activity (1.25 mg/kg) in previous studies (Turner et al., 2015; 2017).

Behavioural testing - assessing effects of SKF 38393 on waiting impulsivity.

Rats had been exposed previously to low doses of PD128907 (0.05, 0.1 or 0.2 mg/kg). After a two-week washout period, SKF 38393 was administered. Fifteen rats were used to investigate the effect of SKF 38393 on 5C-CPT performance using challenging task conditions that consisted of a prolonged ITI (12,14 or 16s) in a one-hour session using a within-subjects design. The prolonged ITIs were presented in random order to counteract the tendency to time responses. Following task acquisition rats received 2, 4 or 6 mg/kg SKF-38393 or vehicle, 30 min before the task. All animals received a normal training session between the testing sessions to ensure performance was maintained at a stable baseline.

Assessing effects of AMPH on waiting impulsivity.

Fifteen rats were used to investigate the effect of AMPH on 5C-CPT performance using challenging task conditions that consisted of a prolonged ITI (8,10 or 12s) in a one-hour session. The prolonged ITIs were presented in random order to counteract the tendency to time responses. Using a within-subjects design trained rats received 0.1, 0.2 or 0.4 mg/kg AMPH or vehicle, 10 min before the task. All animals received a normal training session between the testing sessions to ensure performance was maintained at a stable baseline. Prior to the first test day, all animals had been habituated twice to i.p. saline injections.

### Statistical analysis

All data were displayed as observed mean ± SEM. Overall performance was analysed by one-way repeated measures ANOVA followed by Planned Comparisons. Analysis of performance across the session was carried out by two-way repeated measures ANOVA (Trial and treatment as within-subject factors) followed by Planned Comparison on the predicted means. If the outcome of the repeated-measures ANOVA yielded significant effects of dose, further post-hoc analysis was performed using Dunnett multiple comparison test. Analysis of the performance across ITIs was carried out by two-way repeated measures ANOVA (ITI and Treatment as within-subject factors), followed by Planned Comparisons. Alpha level was set to 0.05 and all analyses were carried out using Prism (v9.0).

### Correlational analysis

This study investigated if baseline performance predicted the degree of change in performance following administration of SKF 38393 or AMPH using Oldham method. Oldham’s method was used to determine the presence of a rate dependent relationship. Pearson’s correlation was used when individuals were normally distributed. Nonparametric Spearman’s correlation was used when individuals were not normally distributed. Oldham’s *r* > 0.3 indicates the presence of a rate dependent relationship (Oldham, 1962; Bickel et al., 2016; Snider et al., 2016; Snider et al., 2018; Abdulkader et al., 2023). The Oldham correlative approach is a more appropriate analysis method when studying compounds with attention and inhibitory response control enhancing potential (Bickle et al., 2016; Abdulkader et al., 2023). The Oldham correlative approach has been used previously to determine the rate-dependent effect of pharmacological manipulation of the dopaminergic system on attention and inhibitory response control (Bickle et al., 2016; Abdulkader et al., 2023). We, therefore, applied correlational analysis in this study to determine if SKF 38393 or AMPH can modulate waiting impulsivity in a baseline-dependent manner.

**Figure 1:**
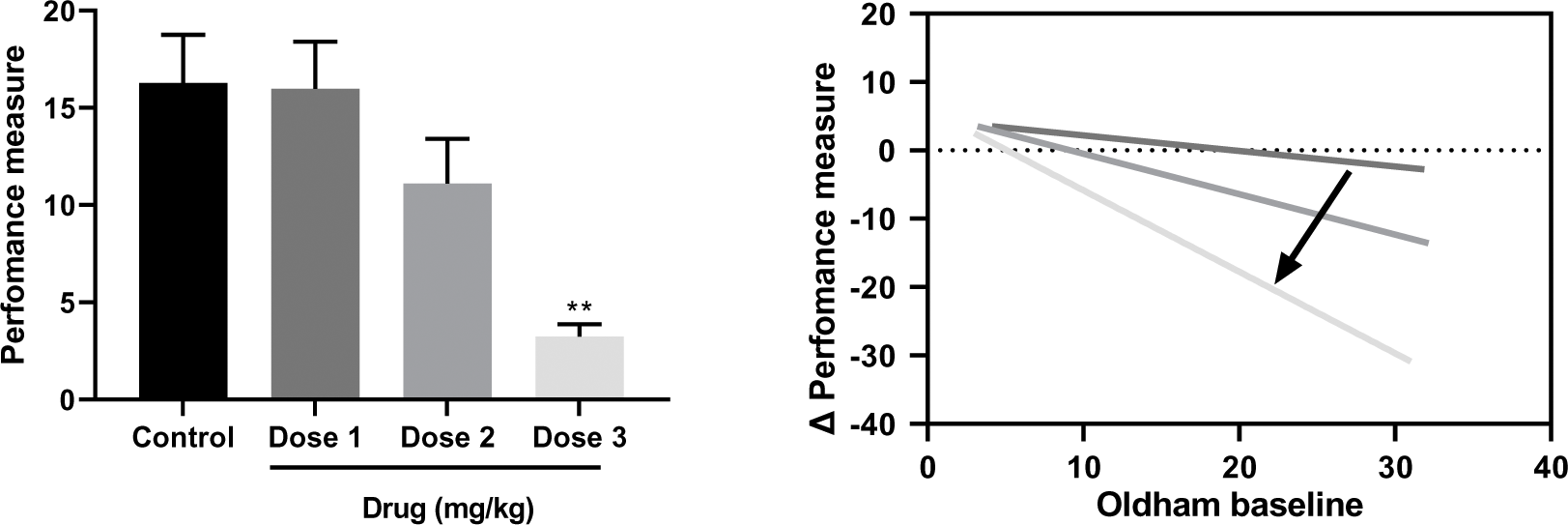
An example of an ideal rate dependent relationship with hypothetical data. The moderate dose shows a tendency towards reducing average performance measure without reaching a significant effect. This is due to a small decrease in high baseline values with no change in the low baseline values. When the highest dose of the drug is administered the decrease in high and low baseline values will produce a significant decrease in the average performance. The clockwise movement of the regression line indicates that, as the dose increases, the magnitude of improvement increases in a manner consistent with baseline performance.

## Results

### Waiting impulsivity

#### Group performance

Effects of SKF-8393 or AMPH on motoric impulsivity is shown in Fig. 2. SKF-38393 produced a significant decrease in 5C-CPT premature responding (F (2.631-31.58) = 6.688, *p* < 0.01; Fig. 2A). Dunnett *post hoc* test revealed a significant decrease in premature responding between vehicle and 6 mg/kg SKF-38393 treated rats (*p*<0.01). AMPH also produced a significant decrease in premature responding in the 5C-CPT (F (1.948, 23.38) = 12.80, *p* < 0.001; Fig. 2B). Dunnett *post hoc* test revealed a significant decrease in premature responding following 0.2 or 0.4 mg/kg compared with control (*p*< 0.001). Effects of SKF 38393 on motoric impulsivity across different ITIs is shown in Fig. 2C; a two-way repeated measures ANOVA revealed a significant treatment effect [F (2.482, 29.78) = 9.686 *p* < 0.001] on impulsivity with no significant ITI effect [F (1.561, 18.73) = 0.1718, P=0.7906] and no interaction [ F (2.673, 32.08) = 0.2205, P=0.8610]. Baseline impulsivity remained stable over all inter-trial intervals. Dunnett’s multiple comparison *post hoc* test revealed that 6 mg/kg SKF 38393 significantly reduced the number of premature responses at 12s (*p*< 0.01), 14s (*p*< 0.05) and 16s (*p*< 0.01) ITIs, indicating that the effect of SKF 38393 was consistent across all intervals. Effects of AMPH on motoric impulsivity over the different inter-trial intervals is shown in Fig. 2D, where a two-way repeated measures ANOVA revealed a significant treatment effect (F [ F (1.970, 23.64) = 12.63, *p* < 0.001) on impulsivity with a significant ITI effect [F (1.286, 15.43) = 9.987, *p* < 0.01] but no interaction [F (2.234, 26.81) = 2.341, P=0.1105]. Baseline impulsivity was significantly higher during 12s compared to 8s ITI. Dunnett’s multiple comparison test revealed that 0.2 mg/kg AMPH significantly reduced the number of premature responses at 8s, 10s and 12s (*p*< 0.05, *p*< 0.05 and *p*< 0.001, respectively) inter-trial intervals, indicating that AMPH effects were consistent across all intervals. Dunnett’s multiple comparison test also revealed that 0.4 mg/kg AMPH significantly reduced motoric impulsivity at 8s (*p*< 0.01), 10s (*p*< 0.01) and 12s (*p*< 0.001) inter-trial intervals, indicating that effect of AMPH was consistent across all inter-trial intervals durations. Effects of SKF on impulsivity across the session is shown in Fig. 2E, where each session consisted of 160 trials broken into 4 blocks, each block consisting of 40 trials. A two-way repeated measures ANOVA revealed a significant effect on impulsivity of trial [F (2.190, 26.28) = 11.26, *p* < 0.001] and treatment [F (2.628, 31.54) = 6.570, *p* < 0.05] as well as a significant interaction [F (4.690, 48.99) = 2.917, *p* < 0.05]. Dunnett *post hoc* tests revealed a significant decrease in the number of baseline premature responses in trial block 4 compared with trial block 1 (*p*< 0.01), indicative of improvement in waiting impulsivity over time. Dunnett tests also revealed that the highest dose of SKF 38393 reduced waiting impulsivity in the first, second and third (*p*< 0.001, *p*< 0.01 and *p*< 0.01, respectively) trial blocks. However, 2mg/kg and 4 mg/kg SKF 38393 reduced waiting impulsivity only in the first trial block (all *p*< 0.05) when compared to saline treated rats. Effects of AMPH across the session are shown in Fig. 2F, where sessions consisted of 160 trials broken into 4 blocks, with each block consisting of 40 trials. A two-way repeated measures ANOVA approach revealed significant trial [F (1.888, 22.66) = 15.72, *p* < 0.0001] and treatment effects [F (1.913, 22.96) = 12.82, *p*< 0.001] on impulsivity and a significant trial period x treatment interaction [F (3.546, 42.55) = 2.748, *p*< 0.05]. Dunnett tests revealed a significant decrease in the number of baseline premature responses in trial block 4 (121-160) compared with trial block 1 (1-40; *p*< 0.01), indicative of improvement in waiting impulsivity over time. Dunnett tests also showed that 0.2 mg/kg and 0.4 mg/kg AMPH reduced waiting impulsivity in all trial periods.

**Figure 2:**
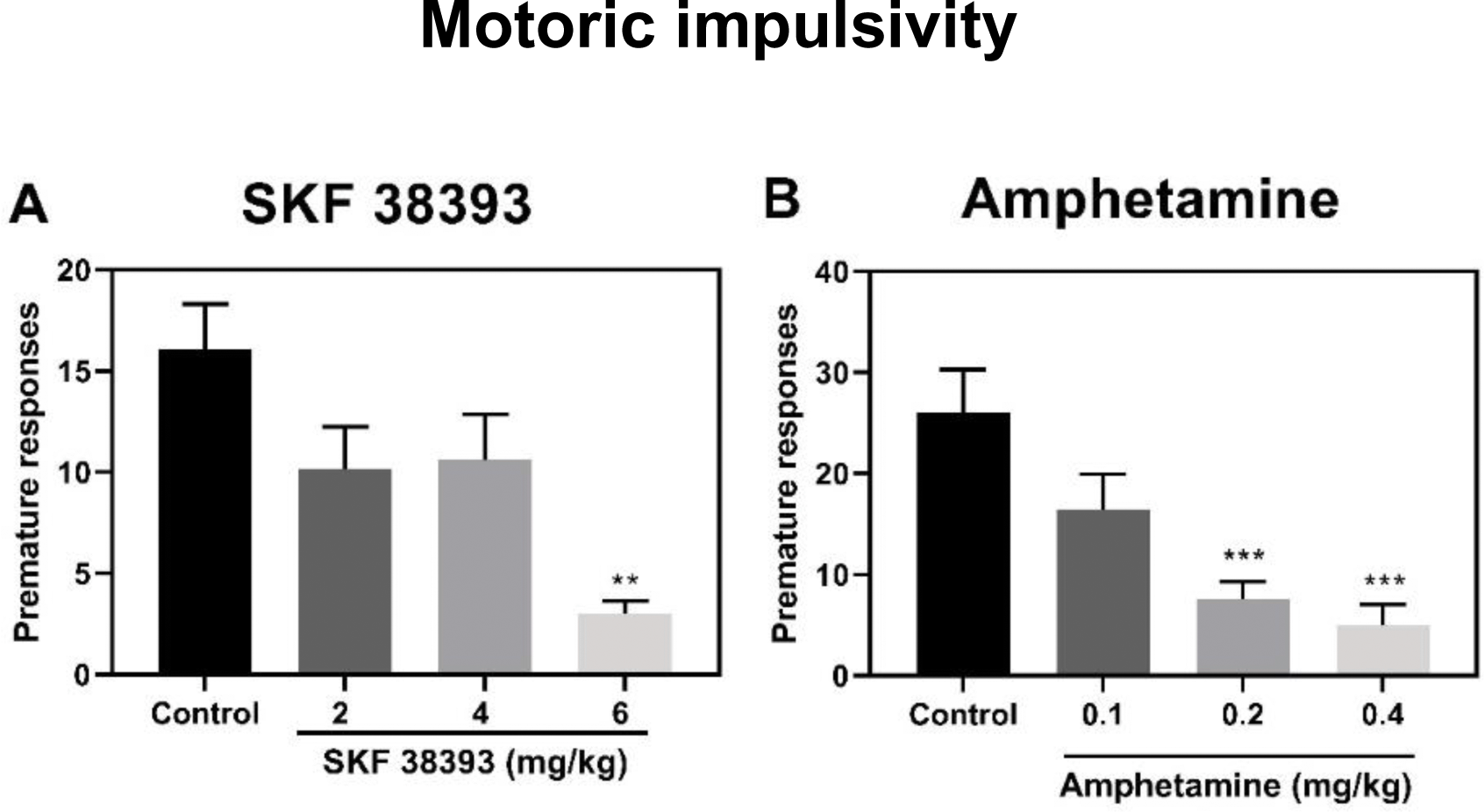
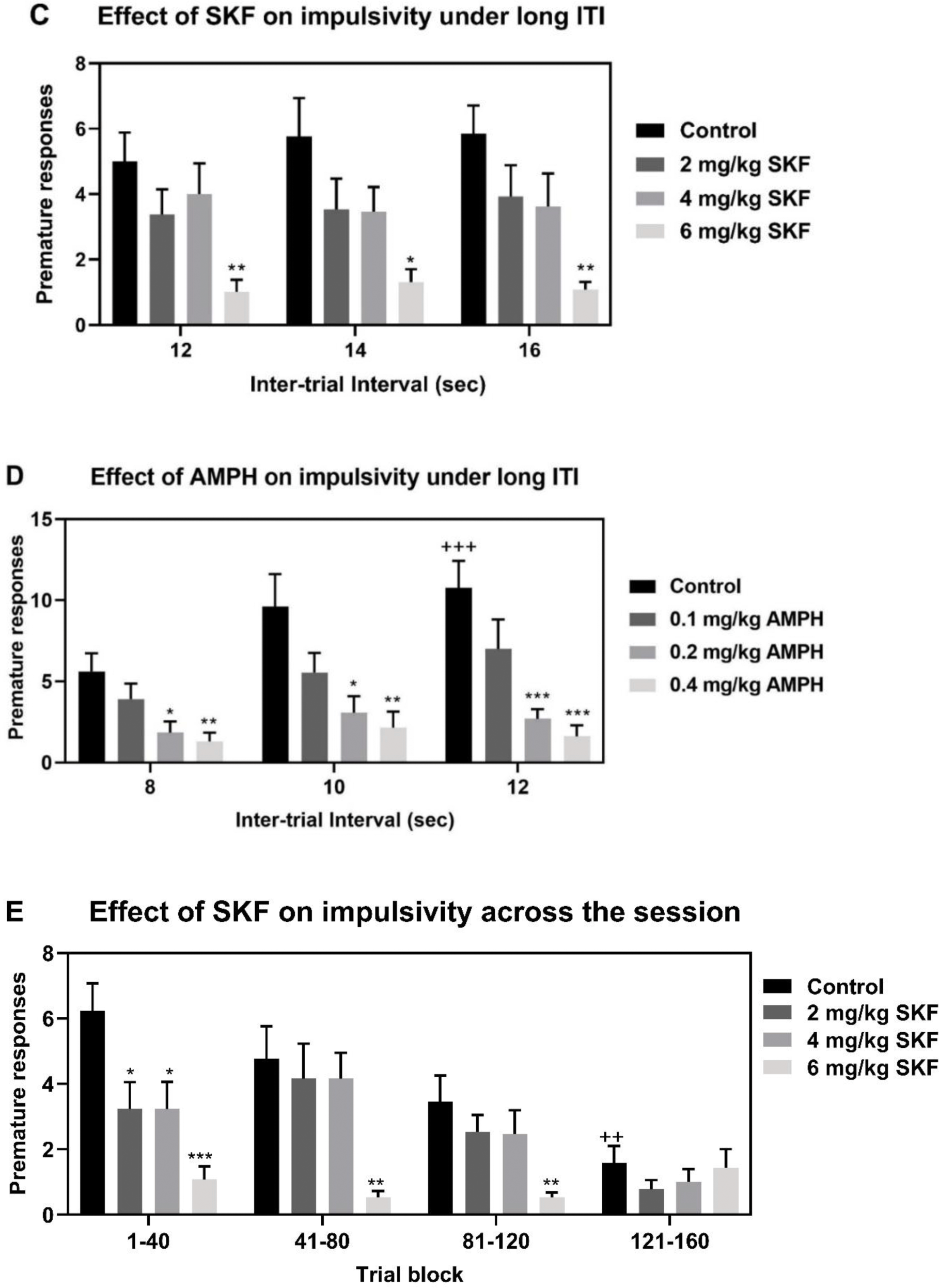
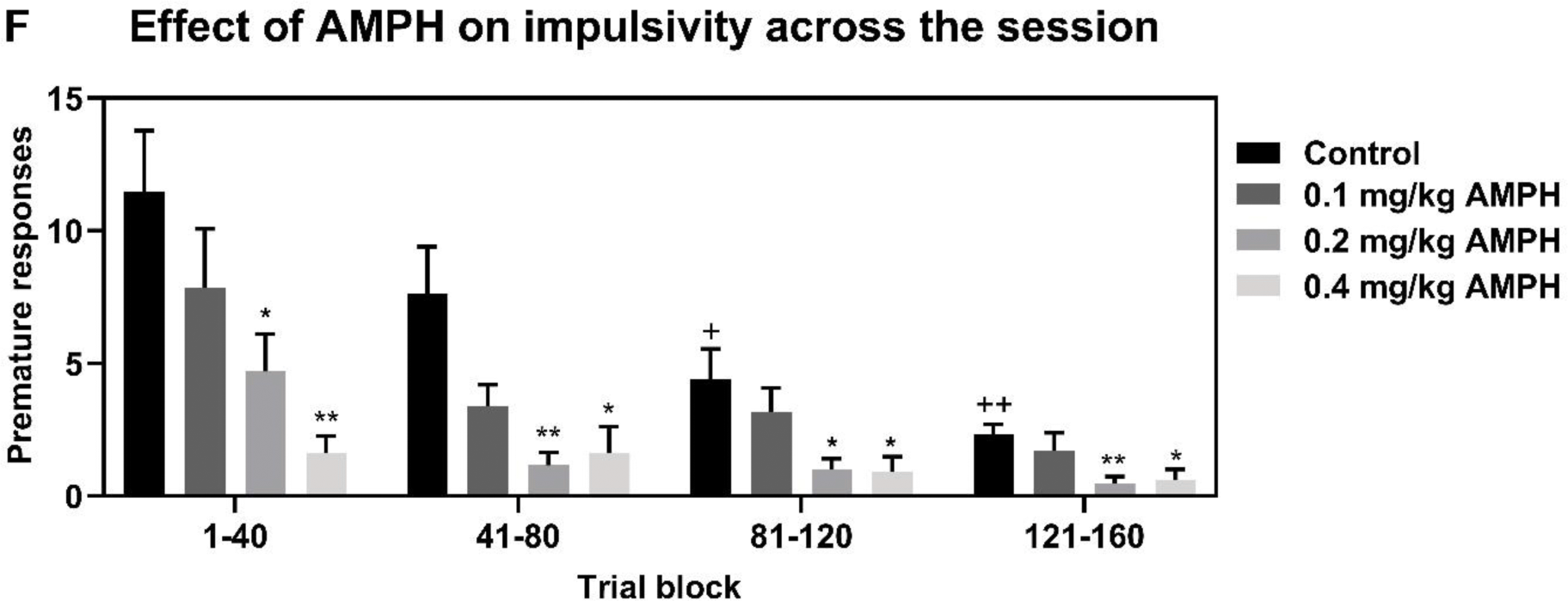
Differential effect of SKF 38393 (2, 4 or 6 mg/kg) or AMPH (0.1, 0.2 or 0.4 mg/kg) on waiting impulsivity of rats tested under long inter-trial intervals. (A and B) Impulsivity averaged over all ITIs. (C and D) Impulsivity over the three different inter-trial intervals. (E and F) Impulsivity over session trial blocks, where each session consisted of 160 trials broken into 4 blocks of 40 trials. * *p* < 0.05, ** *p* < 0.01, *** *p* < 0.001 compared to vehicle-treated rats (Control) of the same ITI or trial block (Dunnett). + *p* < 0.05, ++ *p* < 0.01, +++ *p* < 0.001, ++++ *p* < 0.0001 compared to control of the first block or the shortest ITI (Dunnett).

#### Correlational analysis

The relationship between baseline impulsivity and SKF or AMPH induced change is shown in Fig 3A. There was a significant strong negative correlation between baseline premature response and the SKF-induced change at the highest dose, consistent with rate-dependence. There were also strong negative correlations between baseline premature response and the change following administration of 0.2 mg/kg (r = - 0.78, *p*< 0.01) or 0.4 mg/kg AMPH (r = -0.65, *p* < 0.05), again consistent with rate-dependence. The most impulsive rats showed greater decrease in premature responding following AMPH administration. Oldham regression line can move in clockwise (become steeper) or counterclockwise direction (become more shallow) as the dose increase due to the gradient of the line increasing or decreasing. Here, the Oldham regression line moved in clockwise direction (effectiveness increases). The strong negative inverse relationship between baseline impulsivity and magnitude of improvement following administration of the highest dose of SKF 38393 or the two highest doses of AMPH is responsible for the significant decrease in average performance compared to control.

**Figure 3:**
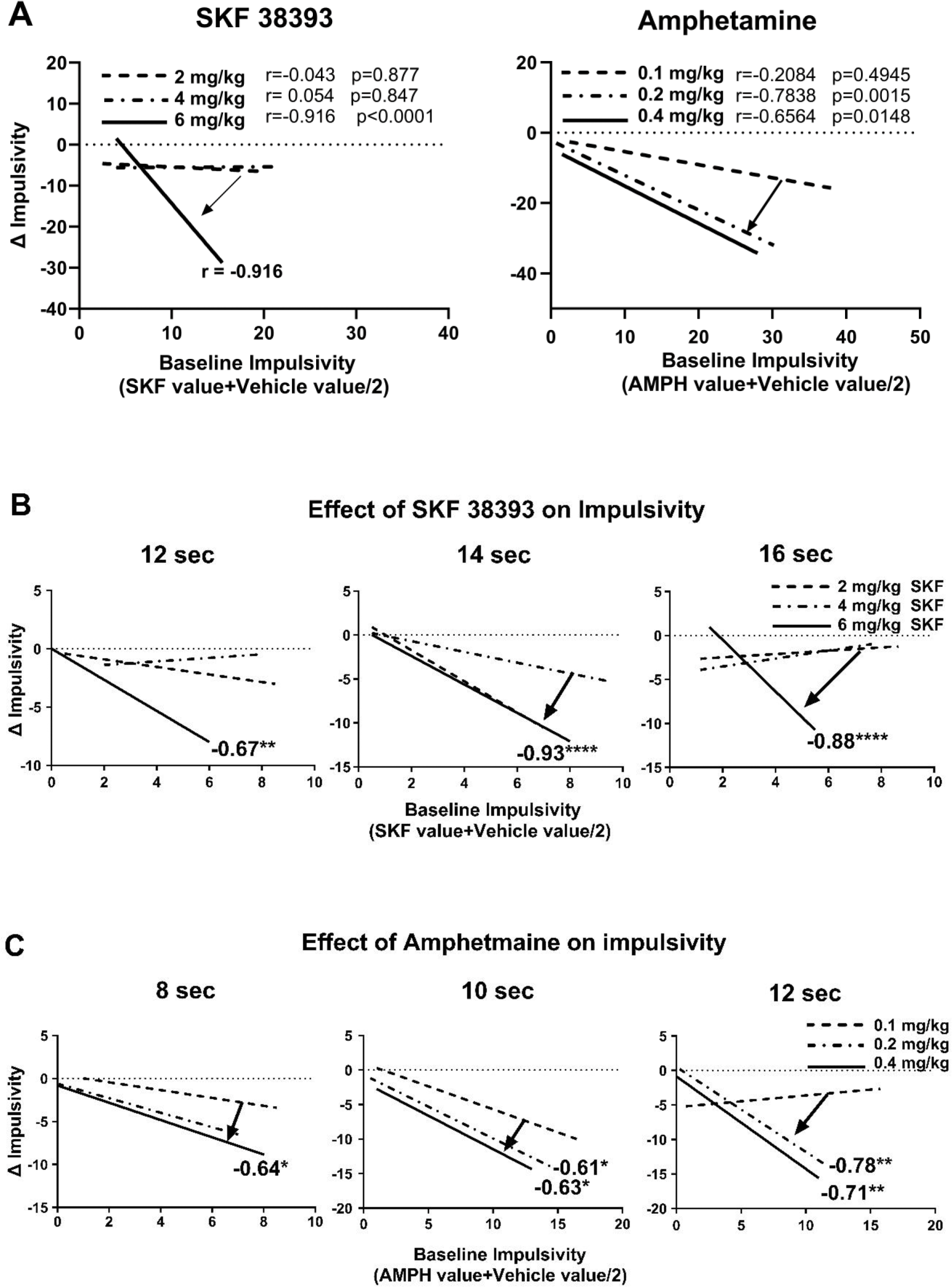
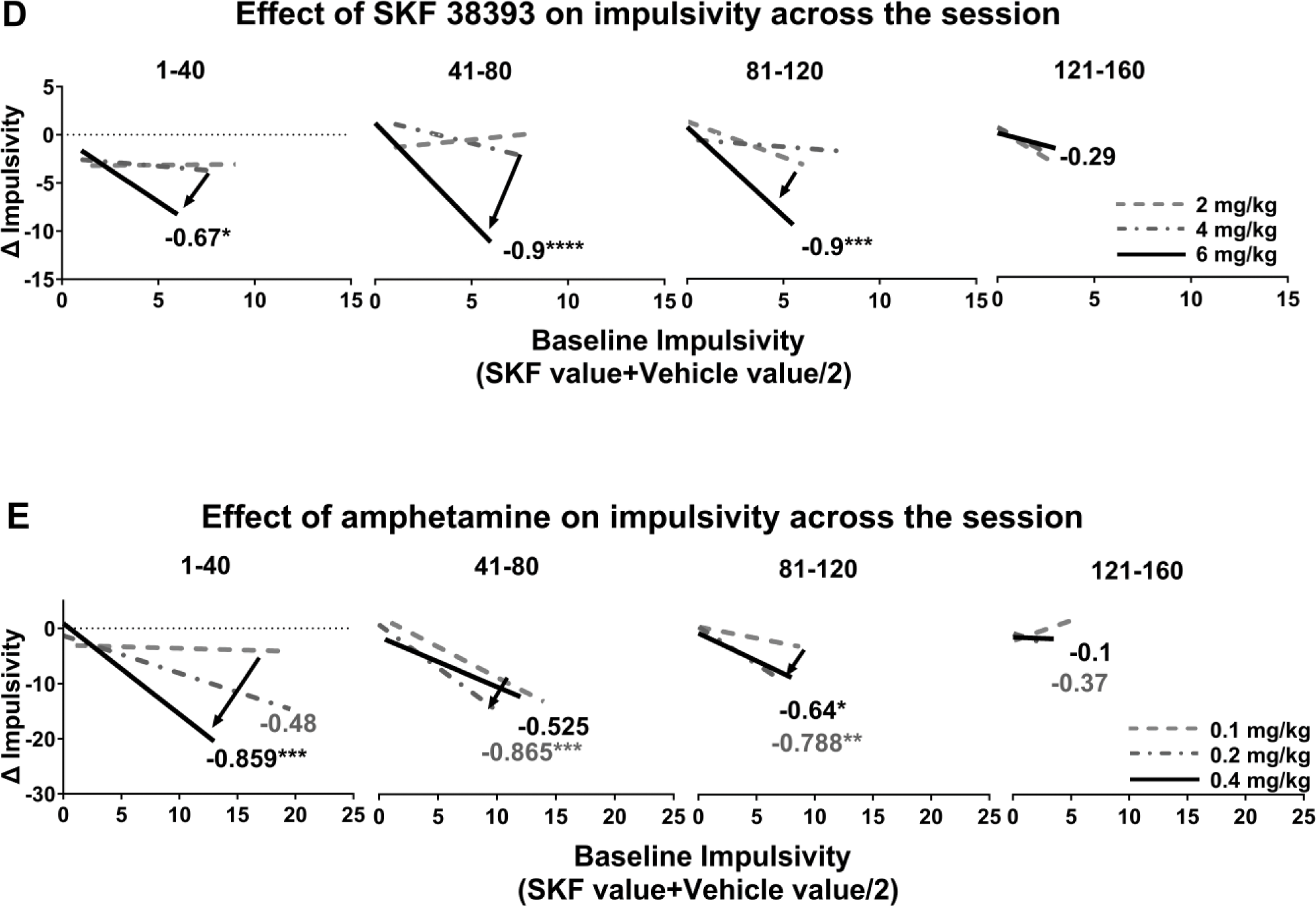
Relationship between baseline impulsivity and SKF 38393 (2, 4 or 6 mg/kg) or AMPH (0.1, 0.2 or 0.4 mg/kg) induced change. (A) Correlation between baseline impulsivity and the drug induced change. (B and C) Correlation between baseline impulsivity and SKF or AMPH-induced change for each ITI. (D and E) Correlation between baseline impulsivity and SKF or AMPH-induced change across the session by trial block. Arrows show the clockwise or counterclockwise movement of the regression line as the dose increases.

As there was wide variability in baseline premature response at each inter-trial interval or trial block, the individual baseline premature responses of rats were normalised by plotting against magnitude of improvement following administration of SKF 38393 or AMPH by ITI or trial block (Fig 3B-E). The highest dose of SKF 38393 or AMPH changed impulsivity in a baseline dependent manner at all ITIs (see Fig 3B-C and Supplement). However, the relationship between baseline impulsivity and magnitude of improvement following administration of the highest dose of SKF 38393 was stronger than the same relationship following administration of the highest dose of AMPH at all ITIs, suggesting that SKF 38393 is more effective than AMPH in this respect. The intermediate dose of AMPH improved waiting impulsivity in a rate-dependent manner for 10s and 12s ITIs (see Fig 3C and Supplement). The highest dose of SKF 38393 improved waiting impulsivity in a baseline dependent manner in the first, second and third trial block (Fig 3D). The relationship between baseline impulsivity and AMPH-induced change across the session is shown in Fig 3D and E. The highest dose of AMPH (0.4 mg/kg) improved waiting impulsivity in the first and third trial block. The intermediate dose of AMPH (0.2 mg/kg) improved waiting impulsivity in the second and third trial block. The highest dose of SKF 38393 was more effective than the highest dose of AMPH in the second and third trial block. The clockwise movement of the regression line as the dose increases is an indicator of improvement in impulsivity.

### Measurements of Oldham’s angles

Angle measurements are shown in Fig 4. Amount of space between the regression lines were measured using a protractor. Three angles were measured: ∠1 (blue), ∠2 (green) and ∠3 (red) as shown in Fig 4. Oldham’s angle measurements for the relationship between baseline impulsivity and SKF 38393 induced change are shown in Fig 4A. ∠1 is the space between 6 mg/kg and 2 mg/kg regression lines. ∠2 is the space between 6 mg/kg and 4 mg/kg regression lines. ∠3 is the space between 4 mg/kg and 2 mg/kg lines (Fig 4A). ∠1and ∠3 are two opposite interior angles. ∠2 is the exterior angle of the triangle (Fig 4A). The Exterior Angle theorem states that the exterior angle of a triangle is equal to the sum of the two opposite interior angles. If two angles are known, then the exterior angle theorem can be used to calculate the third angle. For example, ∠3=∠2-∠1 (Fig 4A). Oldham’s angle measurements for the relationship between baseline impulsivity and AMPH induced change are shown in Fig 4B. ∠1 is the space between the highest and the lowest dose (0.4 mg/kg and 0.1 mg/kg regression lines). ∠2 is the space between the highest and the intermediate dose (0.4 mg/kg and 0.2 mg/kg regression lines). ∠3 is the space between 0.2 mg/kg and 0.1 mg/kg lines (Fig 4B). Line elongation showed that ∠3 and ∠2 are two opposite interior angles. ∠1 is the exterior angle of the triangle (Fig 4B). If two angles are known, then the third angle can be calculated. ∠2=∠1-∠3 (Fig 4B). The clockwise movement (⟳) of the regression line as the dose increases shows that the highest dose is more effective than the lowest or the intermediate dose at reducing impulsivity. The angle measurement and the direction of movement (clockwise or counterclockwise) shows how effective is one dose at lowering impulsivity compared with other doses.

**Figure 4:**
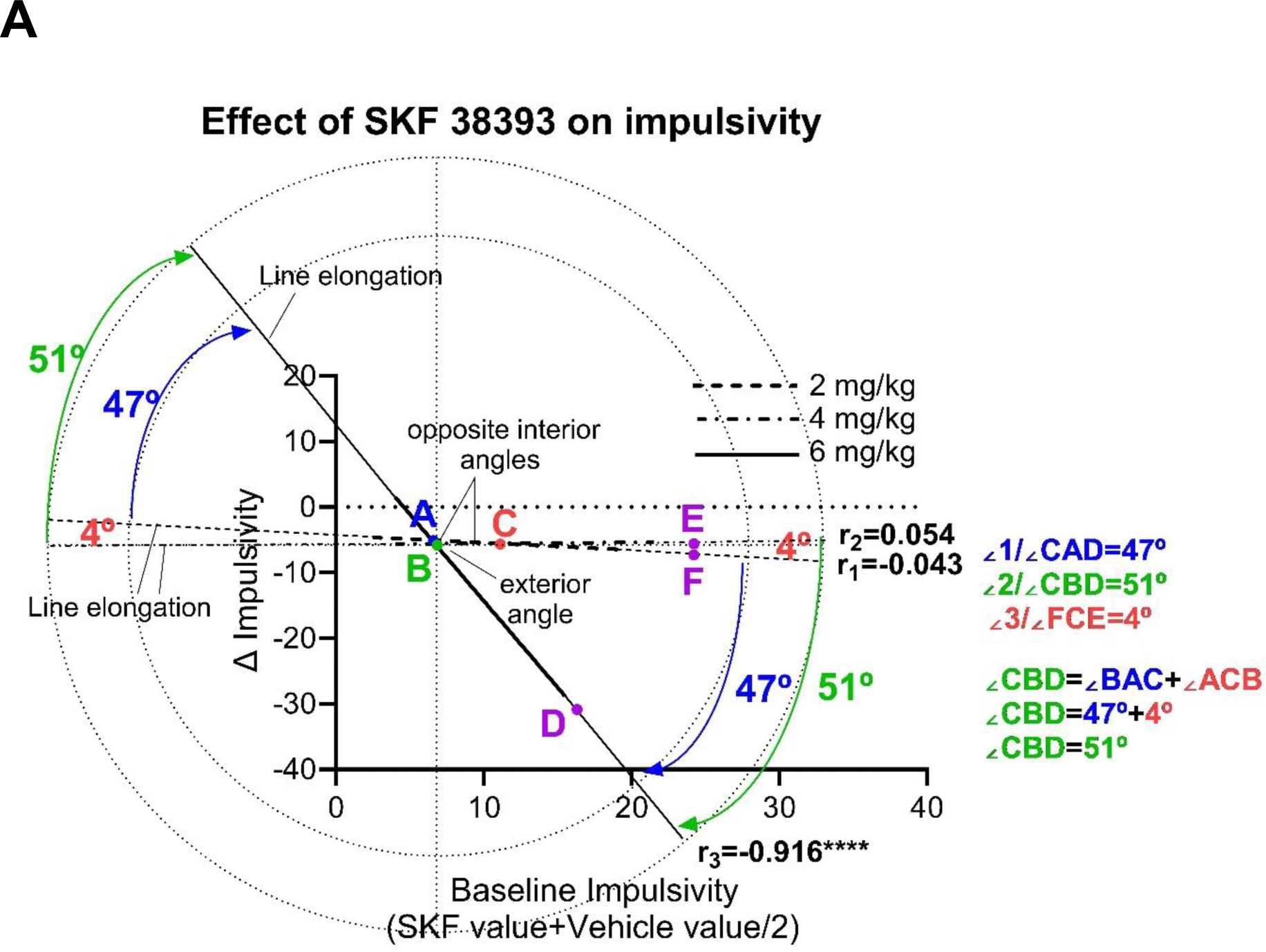
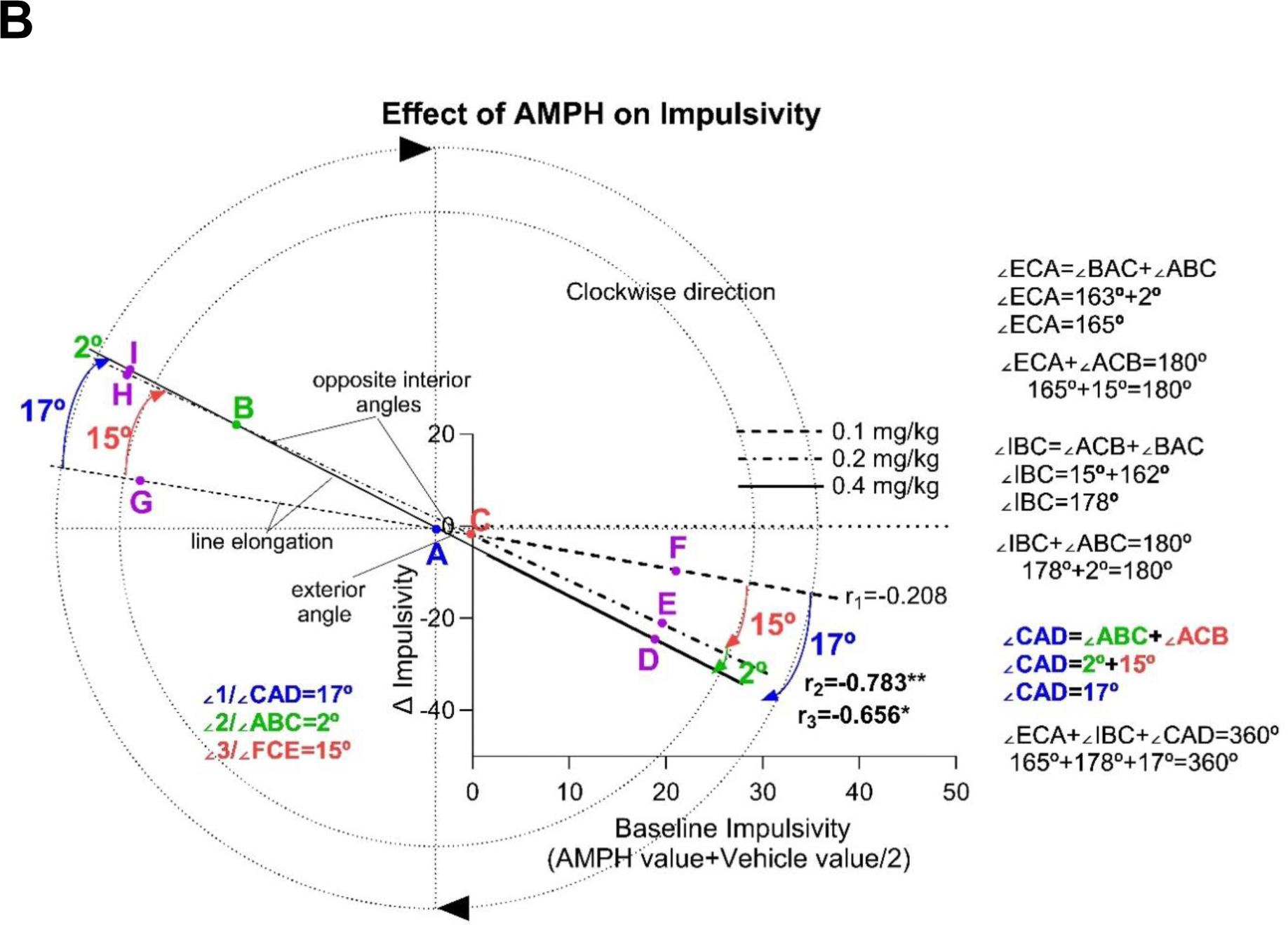
Measurement of Oldham’s angles. (A) Correlation between baseline impulsivity and SKF 38393 induced change. (B) Correlation between baseline impulsivity and AMPH induced change. The Arrows show the clockwise movement of the regression line as the dose increases.

Oldham’s angle measurements for the relationship between baseline impulsivity and SKF 38393 induced change at the shortest ITI are shown in Fig 5A. ∠1and ∠3 are two opposite interior angles. ∠2 is the exterior angle of the triangle. ∠2=∠1+∠3. Oldham’s angle measurements for the relationship between baseline impulsivity and AMPH induced change at the shortest ITI are shown in Fig 5B. ∠2and ∠3 are two opposite interior angles. ∠1 is the exterior angle of the triangle. ∠1=∠2+∠3.

**Figure 5:**
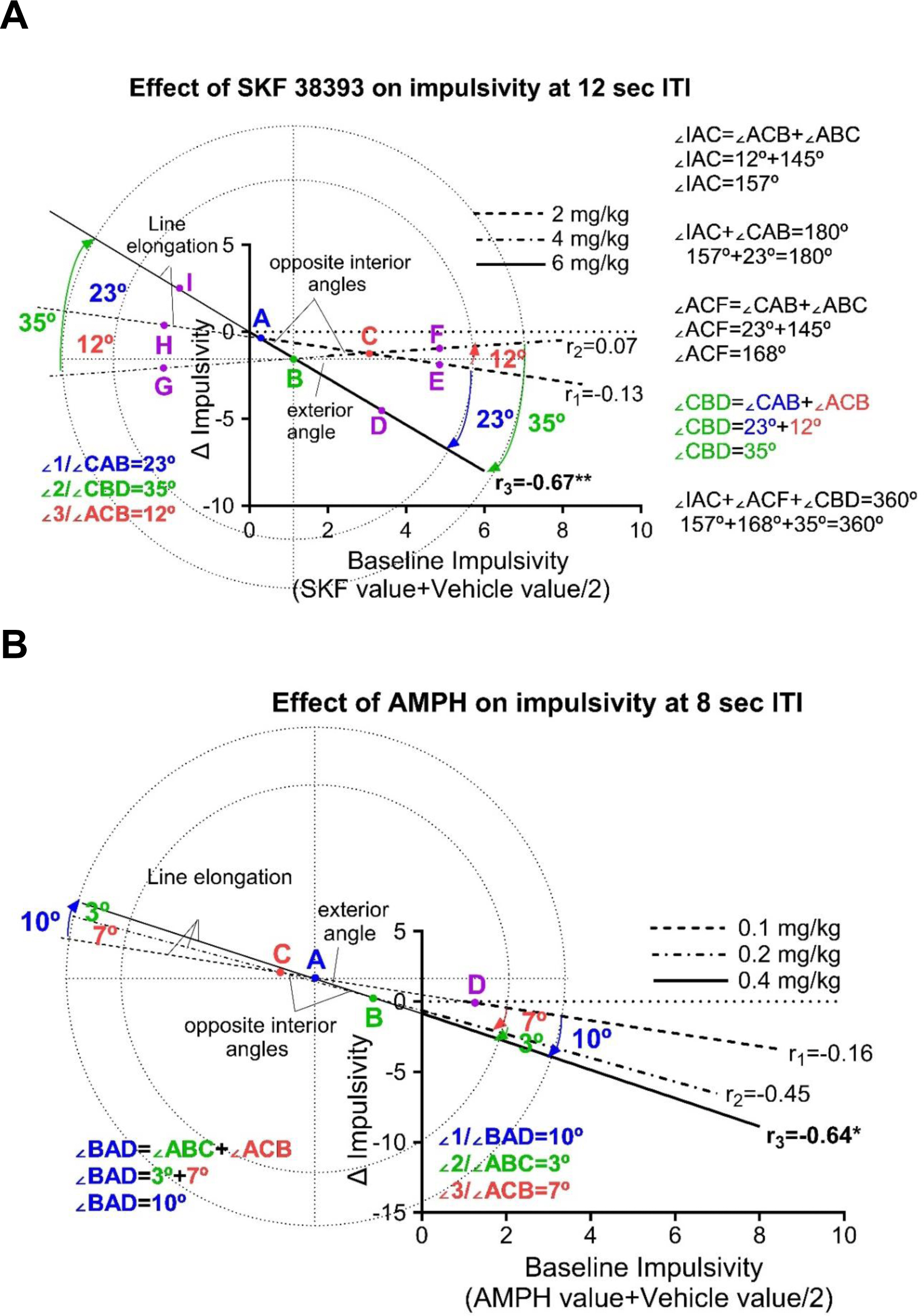
Measurement of Oldham’s angles at the shortest ITI. (A) Correlation between baseline impulsivity and SKF 38393 induced change at 12 sec ITI. (B) Correlation between baseline impulsivity and AMPH induced change at 8 sec ITI.

Angle measurements for the relationship between baseline impulsivity and SKF 38393 or AMPH induced change in the first trial block are shown in Fig 6. ∠2and ∠3 are two opposite interior angles. ∠1 is the exterior angle of the triangle (Fig 6A). ∠1=∠2+∠3 (Fig 6A). ∠2and ∠3 are adjacent angles (Fig 6B). ∠1=∠2+∠3 (Fig 6B)

**Figure 6:**
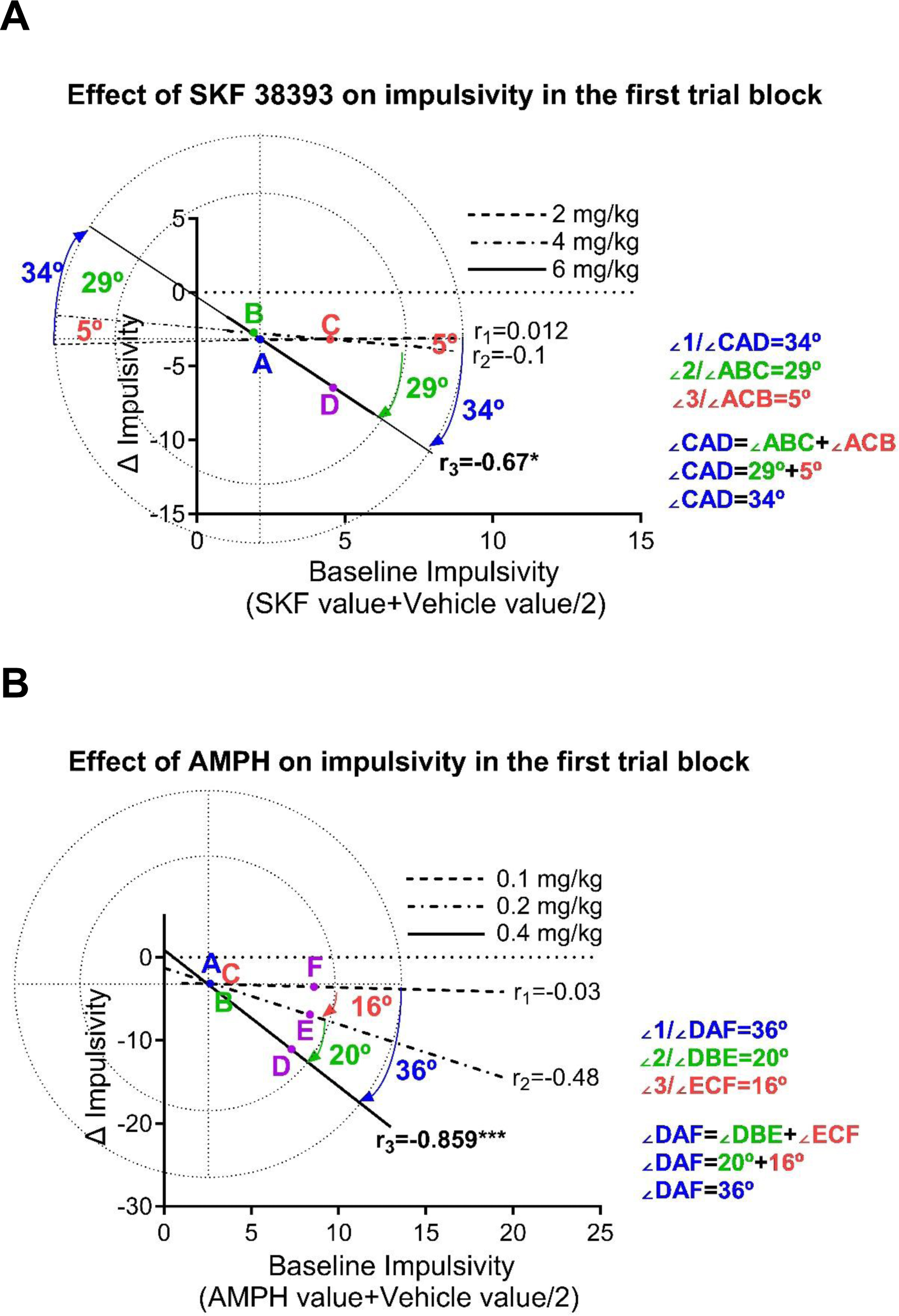
Measurement of Oldham’s angles in the first trial block. (A) Correlation between baseline impulsivity and SKF 38393 induced change. (B) Correlation between baseline impulsivity and AMPH induced change.

Angle measurements for the relationship between baseline impulsivity and SKF 38393 or AMPH induced change in the second trial block are shown in Fig 7. The angle measurement and the direction of movement (clockwise or counterclockwise) shows how effective is one dose at lowering impulsivity compared with other doses.

**Figure 7:**
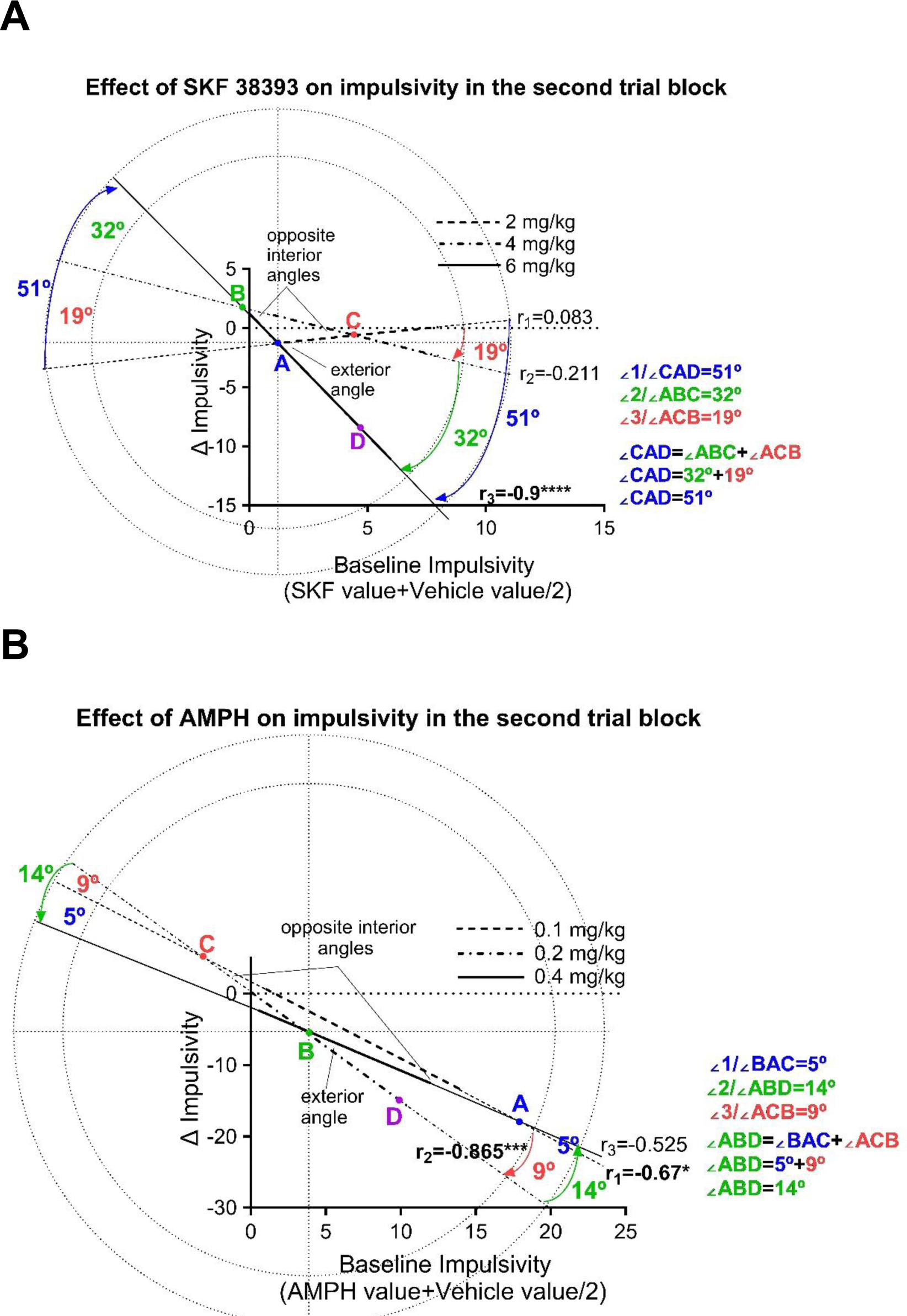
Measurement of Oldham’s angles in the second trial block. (A) Correlation between baseline impulsivity and SKF 38393 induced change. (B) Correlation between baseline impulsivity and AMPH induced change.

Oldham’s angle measurements for the relationship between baseline impulsivity and SKF 38393 or AMPH induced change in the third trial block are shown in Fig 8. The angle measurement and the direction of movement (clockwise or counterclockwise) shows how effective is one dose at lowering impulsivity compared with other doses.

**Figure 8:**
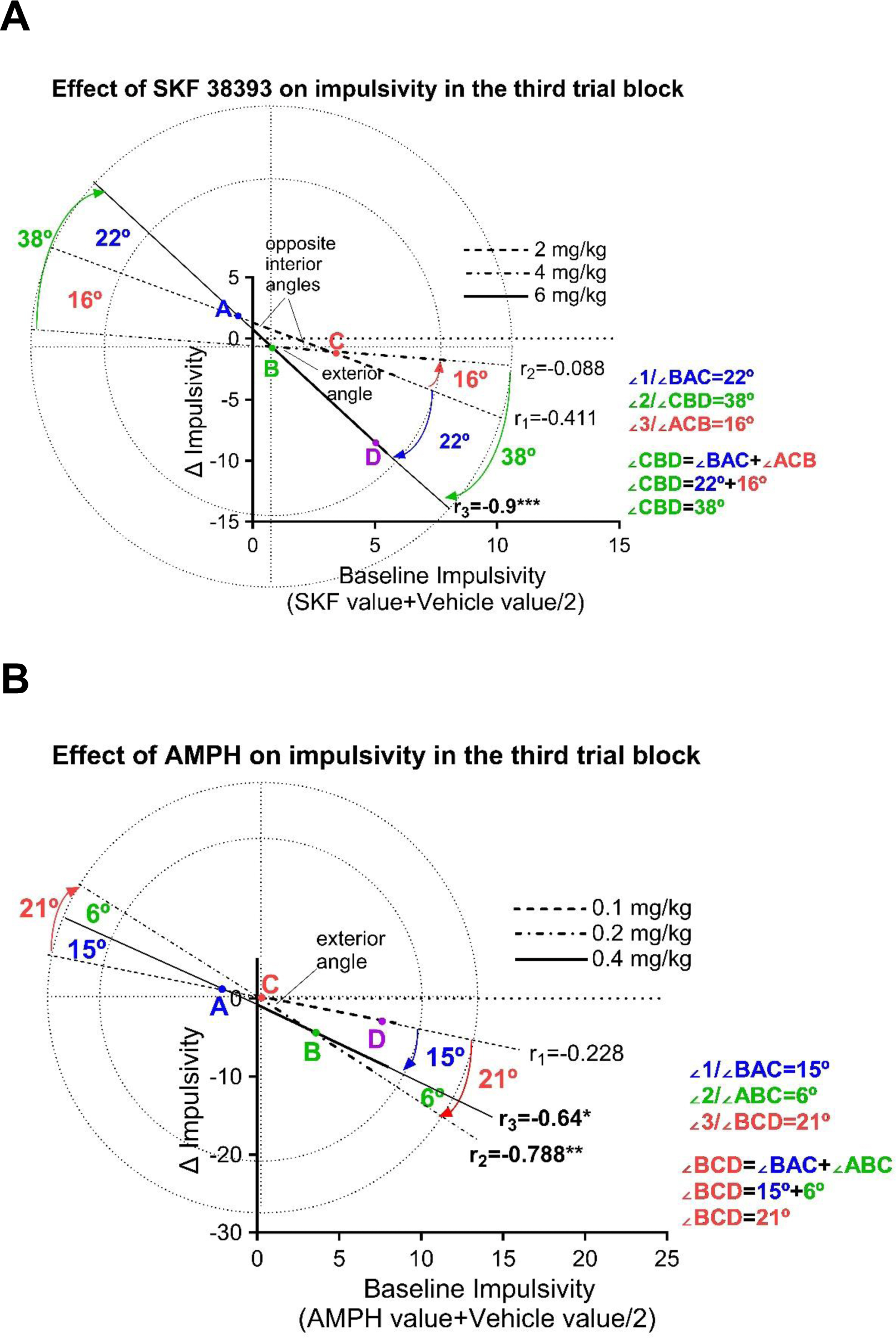
Measurement of Oldham’s angles in the third trial block. (A) Correlation between baseline impulsivity and SKF 38393 induced change. (B) Correlation between baseline impulsivity and AMPH induced change.

### Responsivity index (Motivation)

Effects of SKF on the responsivity index is shown in Fig 9A. A two-way repeated measures ANOVA revealed no significant effect of trial [F (1.430, 17.17) = 0.5885, *p* = 0.5116], a significant treatment effect [F (2.628, 31.54) = 6.570, *p* < 0.05] but no interaction [F (5.153, 52.67) = 3.333, *p* = 0.6538]. There was no significant change in baseline responsivity index (motivation) across the session. Dunnett tests also revealed that the intermediate dose of SKF 38393 increased responsivity in the first and third (*p*< 0.05 and *p*< 0.01 respectively) trial blocks. Effects of AMPH on responsivity index are shown in Fig 9B. A two-way repeated measures ANOVA revealed no significant effect of trial [F (2.392, 31.09) = 1.713, *p* =0.192] treatment [F (1.695, 22.03) = 0.6690, *p* = 0.498] or interaction [F (3.429, 41.52) = 1.726, *p* = 0.17]. Together, these findings suggest that the decrease in baseline waiting impulsivity by the end of the session was mainly due to improvement in inhibitory response control.

**Figure 9:**
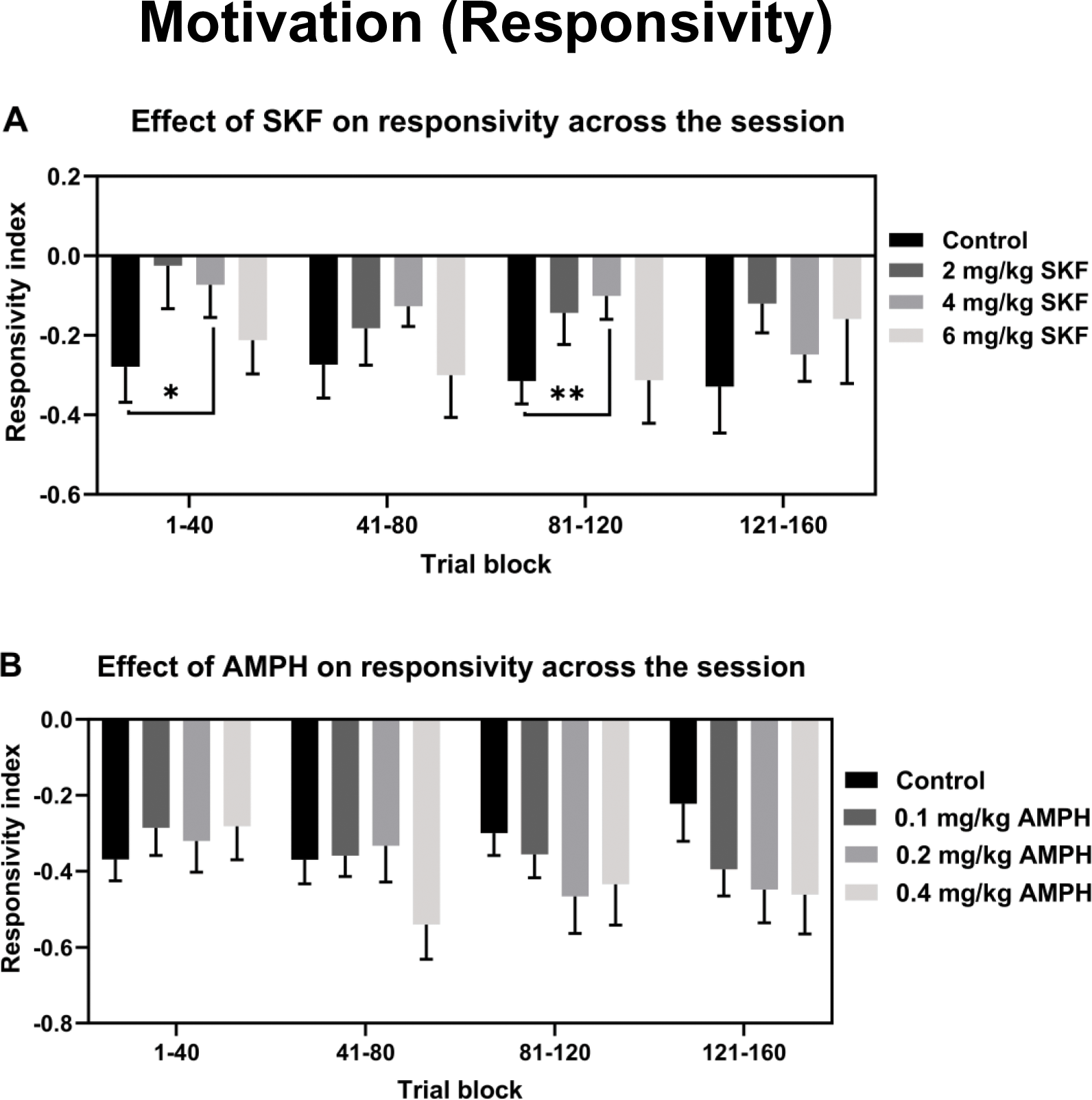
Effects of SKF 38393 or AMPH on responsivity index by session trial block. * *p* < 0.05 and ** *p* < 0.01 compared to vehicle treated rats (Control) of the same block (Dunnett).

### Compulsive behaviour

Effects of SKF 38393 on compulsive behaviour are shown in Fig. 10A. A one-way repeated measures ANOVA revealed no significant effect treatment of [F (1.063, 15.94) = 1.495, P=0.2413] on perseverative correct responses but a significant effect of treatment [F (1.719, 25.79) = 3.617, *p* < 0.05] on perseverative false alarm. Dunnett’s multiple comparison *post hoc* test revealed a significant increase in perseverative false alarm (*p*< 0.05) at 4mg/kg SKF 38393 when compared to control. Effects of AMPH on compulsive behaviour are shown in Fig. 10B. A one-way repeated measures ANOVA revealed no significant effect of treatment [F (1.250, 16.25) = 0.4516, *p* = 0.55] on perseverative correct response or perseverative false alarm [F (2.513, 32.67) = 1.628, *p* =0.2].

**Figure 10:**
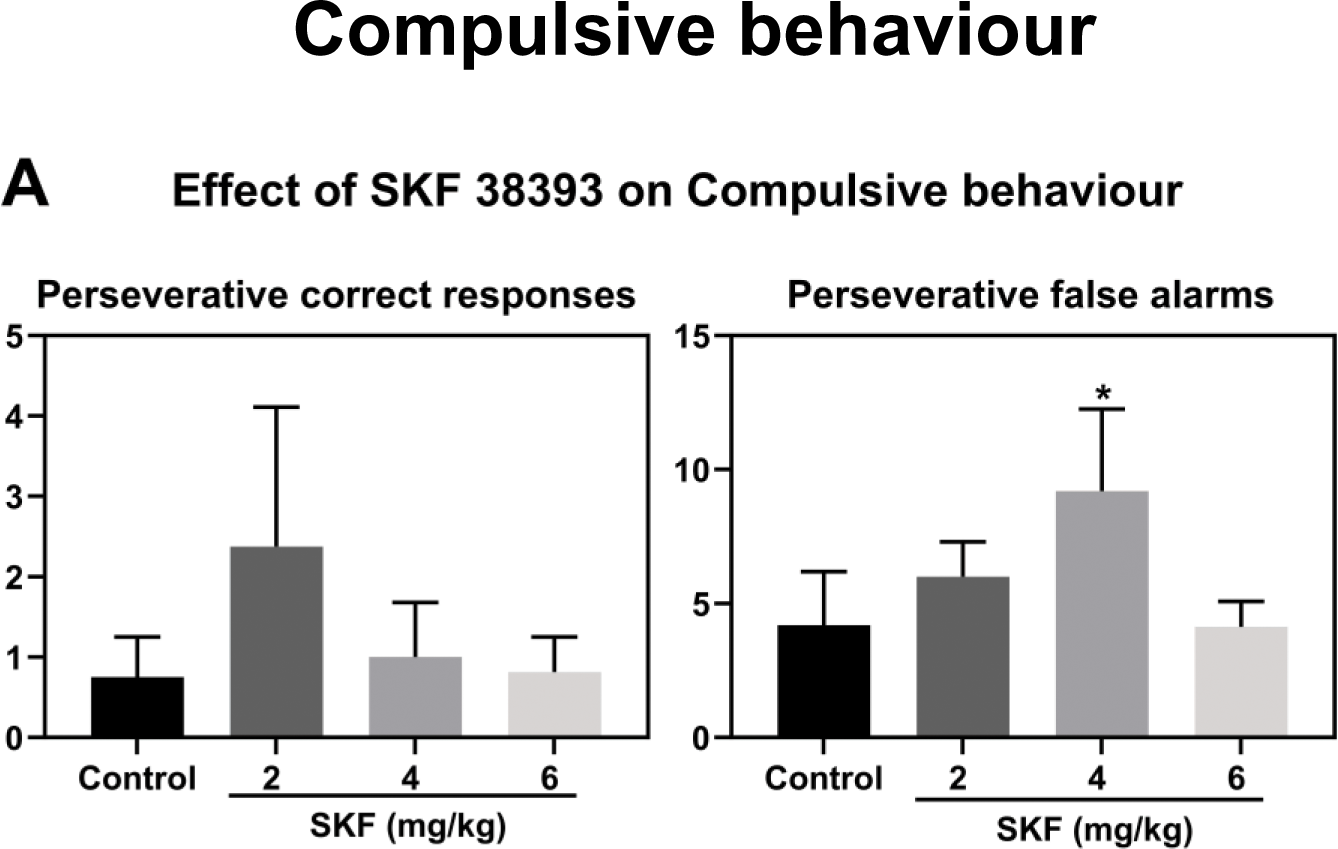
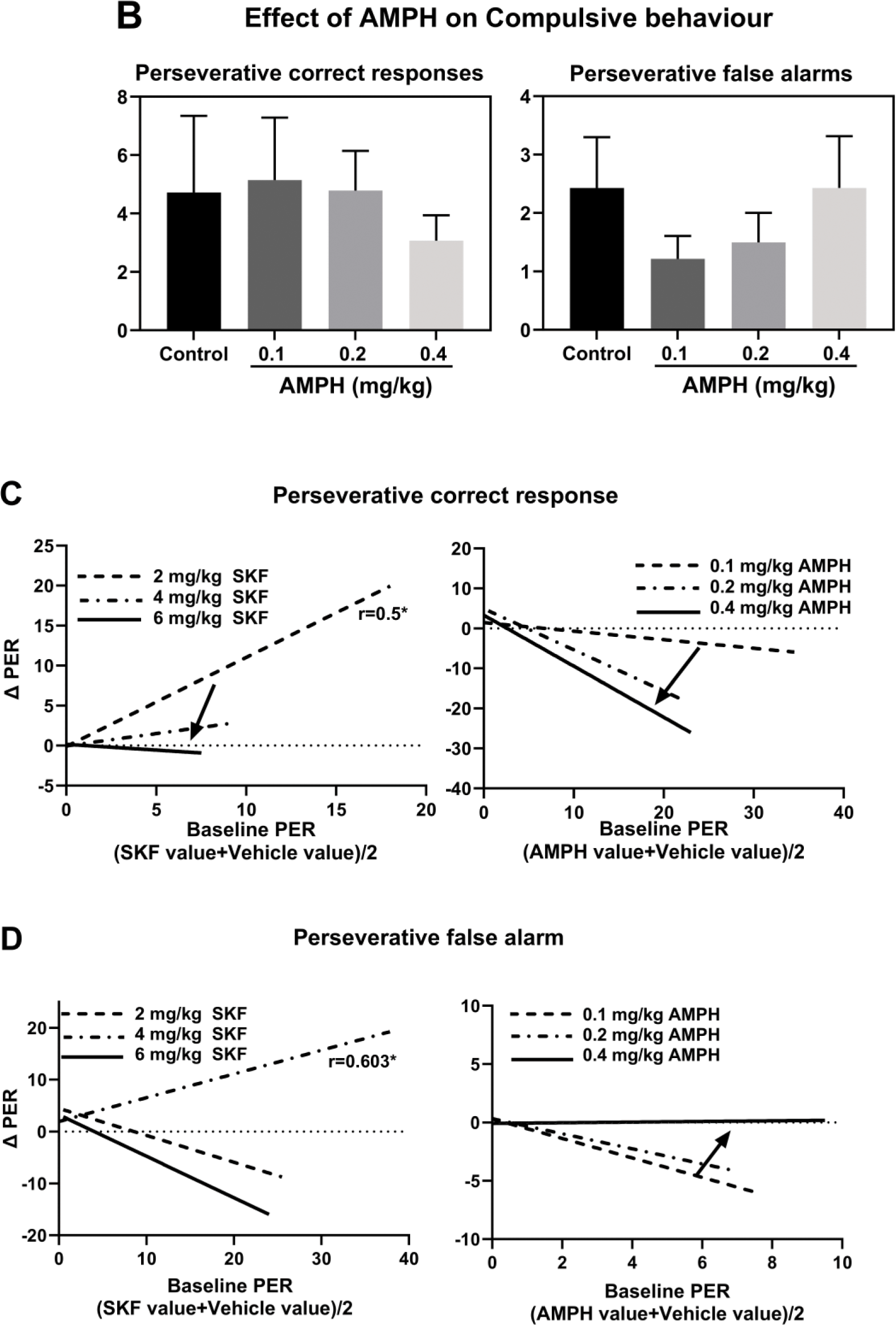
Differential effect of SKF 38393 (2, 4 or 6 mg/kg) or AMPH (0.1, 0.2 or 0.4 mg/kg) on compulsive behaviour of rats tested under long inter-trial intervals. (A and B) Effects of SKF 38393 or AMPH on perseverative correct responses or perseverative false alarms. (C) Correlation between baseline perseverative correct response and SKF or AMPH-induced change. (D) Correlation between baseline perseverative false alarm and SKF or AMPH-induced change. Arrows show the clockwise or counterclockwise movement of the regression line as the dose increases. \**p* < 0.05 compared to vehicle-treated rats (Dunnett). PER = perseverative responses.

The relationship between baseline perseverative response and SKF 38393 or AMPH-induced change is shown in Fig. 10C-D. There was a significant positive correlation between baseline perseverative correct responses and the change following administration of 2 mg/kg SKF 38393 (Fig. 10C), indicative of a rate-dependent effect. In addition, there was a significant positive correlation between baseline perseverative false alarm and the change following administration of 4 mg/kg SKF 38393 (Fig 10D), also indicative of a rate-dependent effect. AMPH did not induce baseline-dependent changes in either perseverative correct responses or perseverative false alarms. The clockwise movement the regression line shows that, when the dose increases, perseverative correct response decreases in a manner consistent with baseline performance. In other words, SKF or AMPH changed perseverative correct responses in a dose-dependent manner (Fig 10C). The counterclockwise movement of the regression line following administration of AMPH shows that perseverative false alarm increases with dose (Fig 10D). Interestingly, AMPH had the opposite effect on perseverative correct responses, suggesting that these two behaviours are independent.

### Compulsive behaviour

Angle measurements for the relationship between baseline perseverative correct response and SKF 38393 or AMPH induced are shown in Fig 11. ∠2and ∠3 are adjacent angles (Fig 11A). ∠1=∠2+∠3 (Fig 11A). ∠2and ∠3 are two opposite interior angles. ∠1 is the exterior angle of the triangle (Fig 11B). ∠1=∠2+∠3 (Fig 11B).

**Figure 11:**
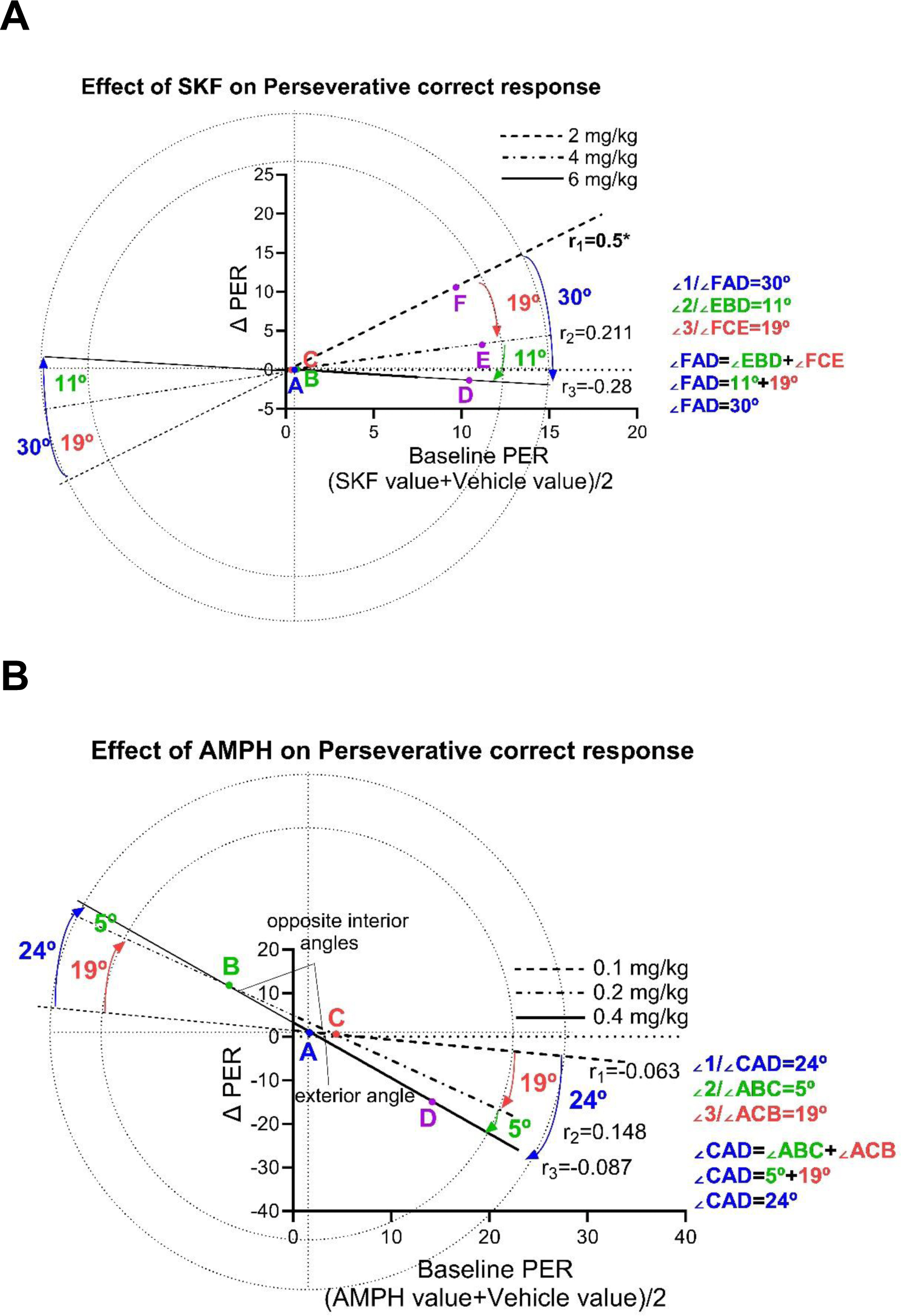
Measurement of Oldham’s angles. (A) Correlation between baseline perseverative correct response and SKF 38393 induced change. (B) Correlation between baseline perseverative correct response and AMPH induced change.

## Discussion

The current study describes a preclinical approach that could help address the urgent need for new ADHD medications that lack the treatment dependence issues seen with currently prescribed medications. The main findings of the present study are: 1) Pharmacological manipulation of dopaminergic system by systemic administration of AMPH or SKF 38393 improves waiting impulsivity in a dose and baseline-dependent manner in female rats; 2) the highest dose of SKF 38393 (6 mg/kg) is more effective than the two highest doses of AMPH in reducing waiting impulsivity; 3) improvement in waiting impulsivity following administration of AMPH verifies predictive validity of the rat version of 5C-CPT; 4) A correlative analysis (i.e., Oldham’s method) is particularly effective in revealing drug effects on behaviour, 5) Analysis of the change in premature responding across the session revealed that the improvement in performance observed following SKF 38393 or AMPH administration is due to enhanced inhibitory response control; and 5) the clockwise movement of the Oldham regression following administration of the highest dose of SKF or the two highest doses of AMPH indicates that, as the dose increases, waiting impulsivity decreases in a manner consistent with baseline performance.

Previous studies suggest that the ability of D1 receptor agonists to improve attentional performance in the 5-CSRTT depends on baseline accuracy performance. For example, systemic administration of SKF-38393 impaired accuracy of rats with high baseline accuracy (Zhu et al., 2017), while direct infusion into the striatum or medial prefrontal cortex increased accuracy of rats with low baseline accuracy (Granon et al., 2000). In addition, previous studies report that methylphenidate or atomoxetine enhances dopaminergic neurotransmission in the prefrontal cortex through indirect stimulation of D1 receptors (Gamo et al., 2010). In the 3-choice serial reaction time task, microinjection of the selective D1 receptor antagonist (SCH 23390) into medial prefrontal cortex blocked the anti-impulsive effects of atomoxetine, suggesting that D1 receptor plays a role in the cognitive enhancing effects of atomoxetine (Sasamori et al., 2019). Furthermore, systemic administration of the D1 receptor agonist SKF 38393 impaired waiting impulsivity in mice when levels of baseline waiting impulsivity were low in 5-CSRTT (Zhu et al., 2017), suggesting that D1 receptor agonists may have beneficial effects on waiting impulsivity under high task demands. However, the role of D1 receptor agonists on waiting impulsivity under high task demand does not seem to have been investigated. The work in this study extends previous findings by investigating the effects of the D1 receptor agonist on waiting impulsivity in the 5C-CPT under challenging task conditions. Previous studies suggest that a prolonged ITI drives animals to make more premature responses, therefore, in this study waiting impulsivity was challenged by extending the ITI. Oldham’s correlation method was used to examine the possibility of improving inhibitory response control as a function of the behavioural baseline. Our findings show that the highest dose of SKF-38393 under high task demand (prolonged ITI) improved waiting impulsivity in a rate-dependent manner. This finding is consistent with pro-attentional effects of 6 mg/kg SKF-38393 (Barnes et al., 2012). This finding is also consistent with previous studies that suggest D1 receptors have an important role in modulating inhibitory response control in 5-CSRTT (Balachandran et al., 2018).

Zhu and colleagues reported worsening in impulsivity following administration of the D1 receptor agonist SKF-38393 under standard task conditions in the 5-CSRTT (Zhu et al., 2017). The finding of the present experiment together with that of Zhu and colleagues suggests that the effectiveness of D1 receptor agonists in improving impulsivity is dependent on baseline activity of the dopaminergic system. In other words, D1 receptor activation worsens impulsivity when the levels of premature responding are low (i.e., under normal task conditions) but improves impulsivity when the levels of premature responding are high. This finding is consistent with the inverted U-shaped relationship between D1 receptor activation and cognitive performance (Arnsten, 1998; Granon et al., 2000). However, our finding contrasts with other studies that report improvement in impulsivity following administration of D1 receptor antagonist (SCH 23390) in the 5-CSRTT (Van Gaalen et al., 2006). It should be noted that the improvement in impulsivity reported by Van Gaalen and colleagues was accompanied by a reduction in the total number of trials completed, therefore, impulsivity improvement could have been secondary to a general decrease in responsivity (Gaalen et al., 2006: Shoaib and Bizarro, 2005). Indeed, other studies have reported that D1 receptor antagonists do not affect waiting impulsivity (Amalric et al., 1993; Zhu et al., 2017). In the present study, the clockwise movement of the Oldham regression line as the dose increased was consistent with performance of rats at group level as the most effective dose of SKF 38393 produced the most significant shift, consistent with previous studies in working memory, which report that highly efficacious treatments improve performance in a rate dependent manner (Bickle et al., 2014). To our knowledge this is the first study to take advantage of the variation in individual baseline waiting impulsivity to demonstrate the presence of an inverse relationship between baseline waiting impulsivity and magnitude of improvement following activation of D1 receptor in rat 5C-CPT. It is clear from these results that the effectiveness of SKF-38393 in improving impulsivity is dependent on dopaminergic system baseline activity. This result also supports the potential value of Oldham’s method in revealing rate-dependent effects of drugs on 5C-CPT. Poor inhibitory response control of high-impulsive rats has been linked with a deficit in dopaminergic function within the NAc (Robinson et al., 2009). In other words, in high-impulsive rats, the dopaminergic system activity is sub-optimal. In this study, the improvement in waiting impulsivity of high-impulsive rats following administration of 6mg/kg SKF 38393 could be due to activation of D1 receptors in the ventral striatum. We can hypothesise that administration of such a D1 receptor agonist increased dopaminergic activity towards a normal level in a manner consistent with baseline performance. Together, these findings suggest that drugs targeting D1 receptors may be useful in improving inhibitory response control in ADHD patients.

Analysis of the change in premature responding across the session revealed a gradual decrease in the number of premature responses, which represents a gradual improvement in waiting impulsivity over time. A within-session learning effect has previously been reported in studies using an extended ITI challenge with a constant stimulus duration (Young et al., 2011). In this study, all doses of SKF-38393 reduced impulsivity at the beginning of the session, however, SKF 38393 failed to improve impulsivity at the end of the session, suggesting that the improvement in performance observed following administration of SKF 38393 is due to enhanced inhibitory response control rather than faster learning of the new challenge. SKF-38393 had no effect on other performance measures such as *d* prime (sustained attention) and accuracy. SKF-38393 only showed improvement in attentional performance in the form of increased hit rate at the intermediate dose (4 mg/kg). However, the highest dose of SKF-38393 did not affect hit rate or false alarm rate. This contrasts with a previous study (Barnes et al., 2012) in which SKF 38393 produced an improvement in sustained attention at 4 and 6 mg/kg in adult rats. However, this difference could be due to the shorter range of ITI used in previous work compared to the present study (8, 9, 10, 11 and 12 s versus 12,14 and 16 s, respectively) (Barnes et al., 2012). Therefore, the challenging conditions under which the study is conducted can be a determining feature of whether the drug improves or impairs performance.

Compulsive behaviour in the 5C-CPT can be defined as the number of repetitive nose pokes following either a correct response or a false alarm (Young et al., 2013). The Oldham method revealed a positive relationship between baseline perseverative correct responses and magnitude of change following administration of the lowest dose of SKF 38393. Despite the lowest dose of SKF 38393 (2 mg/kg) changing perseverative correct responses in female rats in a rate dependent manner, perseverative false alarms remained unaffected in these rats; conversely, the intermediate dose of SKF 38393 (4 mg/kg) changed perseverative false alarms, but not perseverative correct responses, in female rats in a rate-dependent manner. The presence of non-target trials in 5C-CPT provides the ability to assess sustained attention and responsivity index (not possible in the 5-CSRTT). The two lowest doses of SKF-38393 increased responding in general (increased hit rate and false alarm rate) and increased compulsive behaviour in the absence of any effect on waiting impulsivity. On the other hand, the improvement in performance following administration of 6mg/kg SKF 38393 appears to be specific to waiting impulsivity. In other words, the highest dose of SKF 38393 did not affect responsivity index and compulsive behaviour. It should be noted that the increase in the number of perseverative correct responses or perseverative false alarms following administration of SKF 389393 (2mg/kg or 4 mg/kg) reported in this study was accompanied by increase in responsivity index (motivation), therefore, the change in compulsive behaviour could have been secondary to a change in motivation (Gaalen et al., 2006: Shoaib and Bizarro, 2005). Indeed, other studies have reported that the D1 receptor agonist SKF 38393 does not affect perseverative responses as measured by 5-CSRTT (Pezze et al., 2007).

AMPH is an indirect catecholaminergic receptor agonist, increasing extracellular levels of dopamine, noradrenaline and serotonin by four mechanisms: (a) inhibiting reuptake of monoamines, (b) reversing monoamine transporter action, (c) increasing the release of monoamines from their storage vesicles and (d) inhibiting monoamine oxidase, the enzyme responsible for monoamine breakdown (Calipari et al., 2013). Previous studies have shown that individual variability in baseline performance can predict AMPH effects in behavioural pharmacology studies (Perkins, 1999). However, unlike Oldham’s approach, the correlation method traditionally used to examine baseline-dependency is affected by mathematical coupling and regression to the mean (Bickel et al., 2016; Snider et al., 2016). Therefore, in some studies, effects related to individual variability are often ignored by either: (a) using average performance (group means); or (b) dividing animals into subgroups of high and low performers (calculated from baseline level of performance) to examine effects of pharmacological agents on each subgroup (Tomlinson et al., 2014; Hayward et al., 2016; Caballero-Puntiverio et al., 2020). Dividing rats into sub-groups has recently received a great deal of attention in the literature; for example, systemic administration of AMPH (0.3 mg/kg) improved vigilance when baseline vigilance was low in mouse touchscreen 5C-CPT (MacQueen et al., 2018). In addition, AMPH improved vigilance of mice exhibiting low baseline response (about 0.7 d prime) without having a significant effect on mice exhibiting high baseline response (about 1.3 d prime). In the same study, AMPH improved impulsivity of high-impulsive mice but had little effect on impulsivity in low-impulsive mice (Caballero-Puntiverio et al., 2019). Our findings show that administration of AMPH, an indirect dopaminergic agonist, under high task demand (prolonged ITI) improved waiting impulsivity in a rate-dependent manner without affecting attention or response inhibition. In addition, Oldham’s method revealed a limited high-range effect on baseline impulsivity; high baseline values decreased with no change in low baseline values. This finding is consistent with previous studies in the 5-CSRTT that report improvement in waiting impulsivity of rats following the administration of AMPH (0.3-0.9) mg/kg (Bizarro et al., 2004; Bizarro and Stolerman, 2003). However, it is in contrast with other studies in mice that report no change in waiting impulsivity (tested under a prolonged ITI with a mean of 5 s) following administration of AMPH (0.3-1) mg/kg (Yan et al., 2011). Re-analysis of the data from Yan et al. (2011) using Oldham’s method demonstrated the presence of a significant moderate correlation between baseline impulsivity and the change following 1 mg/kg AMPH (Oldham’s *r* =0.5622*). Distribution of the data was consistent with a full range effect. Low baseline values increased while high values decreased following AMPH administration, suggesting that the increase in low baseline values concealed the decrease in high baseline values when the data were analysed in the traditional way (i.e., as one group) (Bickel et al., 2016). Consistent with Yan et al. (2011), AMPH (0.25 and 0.5 mg/kg) did not affect premature responding of mice tested under a fixed ITI (5-sec) in the 5-CSRTT (Fletcher et al., 2013). Reanalysis of the data using Oldham’s method revealed a low range effect consistent with rate dependence. Low baseline values increased with no change in the high baseline values (Bickel et al., 2016). This study is consistent with previous clinical studies which report that AMPH and methylphenidate reduce impulsivity in individuals with high baseline impulsivity (Arkell et al., 2022). The rate-dependent effect of AMPH on inhibitory response control was consistent with effects of 6 mg/kg SKF-38393, suggesting that AMPH modulates inhibitory response control partly through noradrenaline transporters and D1 receptors, most likely within the NAc. This finding is consistent with previous studies reporting that D1 receptors play an important role in the cognitive enhancing effects of stimulants (Arnsten and Dudley, 2005). These findings suggest that it is vital to consider correlational analysis when analysing 5C-CPT data. This approach has allowed us to show here that AMPH reduces impulsive action and that the effect of AMPH on waiting impulsivity was dose- and baseline-dependent. In addition, AMPH improved waiting impulsivity under circumstances when task performance was sub-optimal, suggesting that the dopaminergic system modulates performance under demanding circumstances. This finding further supports the predictive validity of the 5C-CPT and the ability of the 5C-CPT to distinguish inhibitory response control elements. Together, these findings suggest that systemic administration of the D1 receptor agonist or AMPH improves waiting impulsivity in a rate-dependent manner. Further work is required to assess effects of AMPH on other forms of inhibitory response control, such as response inhibition, using Oldham’s method.

In summary, this study has shown an inverse relationship between baseline impulsivity and magnitude of change following administration of either the highest dose of SKF 38393 or the two highest doses of AMPH, further supporting the potential value of the Oldham method in revealing rate-dependent effects of drugs in the 5C-CPT. This study is the first demonstration of enhanced inhibitory response control (waiting impulsivity) after selective activation of D1 receptors, suggesting that D1-selective agonists may have clinical utility for the treatment of impulsive behaviour related disorders such as ADHD. In addition, D1-selective agonists could be even more effective than AMPH. Effects of AMPH on waiting impulsivity further support the predictive validity of the 5C-CPT for assessing pharmacologically-induced improvement in waiting impulsivity. Correlational analysis using the Oldham method may improve our understanding of the neurochemical basis of ADHD and may help to develop new therapeutic agents.

